# Molecular Mechanisms Underlying the Regulation of VCAM-1 Expression by the Short-Chain Fatty Acid Butyrate

**DOI:** 10.64898/2025.12.15.694447

**Authors:** Vrishali S. Salian, Vaishnavi Veerareddy, Xiaojia Tang, Yang Xiao, Krishna R. Kalari, Purna C. Kashyap, Karunya K. Kandimalla

**Author notes:** **Corresponding author** Name: Karunya K. Kandimalla.

## Abstract

Over the past decade, cerebrovascular inflammation has been increasingly recognized as a contributor to the progression of neurodegenerative diseases, particularly Alzheimer’s disease (AD). One of the molecular hallmarks of cerebrovascular inflammation is the increased expression of vascular cell adhesion molecule (VCAM)-1 on blood-brain barrier (BBB) endothelial cells. Exposure to amyloid beta (Aβ) peptides, one of the primary hallmarks of AD, and pro-inflammatory cytokines such as tumor necrosis factor-alpha (TNF-α) induces VCAM-1 expression on the BBB endothelium, which facilitates extravasation of leukocytes into the brain thereby promoting an inflammatory response. Therefore, it is crucial to explore therapeutic agents that can inhibit VCAM-1 expression induced by Aβ and TNF-α. Short-chain fatty acids, such as butyrate, produced by the gut microbiota as byproducts of dietary fiber metabolism, are recognized for their anti-inflammatory properties. In this study, we successfully tested the hypothesis that butyrate mitigates Aβ and TNF-α-induced VCAM-1 expression in polarized human cerebral microvascular endothelial cell monolayers, a widely used BBB in vitro model. Our findings indicated that pre-treatment with butyrate significantly reduced Aβ42 and TNF-α mediated upregulation of VCAM-1. Furthermore, we have shown STAT3/GATA6 axis as a key mediator of anti-inflammatory effects of butyrate. These findings provide mechanistic insight into butyrate’s protective role and highlight its potential to mitigate Aβ and TNF-α-induced cerebrovascular inflammation in AD.

## INTRODUCTION

In Alzheimer’s disease (AD) patients, increased exposure to toxic Aβ peptides, augments cerebrovascular inflammation by increasing the expression of adhesion molecules such as vascular cell adhesion molecule 1 (VCAM-1) and intracellular cell adhesion molecule 1 (ICAM-1). An increase in the expression of these adhesion molecules, particularly VCAM-1, promotes the trans endothelial migration of leukocytes across the BBB endothelium[10, 11]. Upon traversing the BBB, these leukocytes can activate resident immune cells such as astrocytes and microglia, which in turn may trigger the release of pro-inflammatory cytokines such as tumor necrosis factor (TNF)-alpha(α)[12]. Furthermore, studies demonstrated that TNF-α, upon binding to the TNF receptor can activate the mitogen-activated protein kinase (MAPK) signaling pathway, which further increases the expression of VCAM-1[13].

Previous reports have indicated that Aβ exposure can also trigger cerebrovascular inflammation by stimulating the release of TNF-α through the direct activation of microglia and astrocytes. Alternatively, Aβ could induce oxidative stress in BBB endothelial cells, resulting in the release of reactive oxidative species that further augment TNF-α release[14, 15]. Thus, Aβ exposure to the BBB endothelium triggers a perpetual cycle of inflammation between inflammatory cytokines and adhesion molecules. This underscores the importance of investigating and monitoring changes in the expression of these adhesion molecules, and explore therapies aimed at mitigating cerebrovascular inflammation.

The gut microbiome produces bioactive compounds such as short-chain fatty acids (SCFAs), mainly consisting of butyrate, propionate, and acetate that can modulate brain function via gut-brain axis[16–18]. Studies have demonstrated that SCFAs such as butyrate can significantly reduce amyloid burden in AD transgenic mice (APP/PS1)[19, 20]. However, in AD patients a reduction in butyrate levels have been noted[21–23], which could potentially increase Aβ burden and exacerbate cerebrovascular inflammation. Butyrate mediates its physiological effects through either G protein coupled receptors or histone deacetylases (HDACs), which are responsible for their anti-inflammatory properties[24]. Previous studies have demonstrated that butyrate reduces the TNF-α mediated increase in VCAM-1 expression due to its HDAC inhibitory effects[25].

Although these findings provide evidence of the effects of butyrate on VCAM-1 expression, the mechanisms underlying this phenomenon are not well understood. Similarly, mechanistic relationship between butyrate and Aβ/TNF-α exposure on the expression of adhesion molecules in BBB endothelial cells has only been partially explored. In the current study, we tested the hypothesis that butyrate downregulates Aβ42 or TNF-α induced increase in VCAM-1 expression, which is critical to regulate cerebrovascular inflammation. Furthermore, we investigated signaling changes in the BBB endothelium that drive this phenomenon.

## RESULTS

Prior research has elucidated molecular mechanisms that govern the expression of cerebrovascular adhesion molecules, VCAM-1 and ICAM-1, upon exposure to Aβ peptides[26, 27] or pro-inflammatory cytokines[28]. Most of the published evidence highlights the role of the MAPK and nuclear factor kappa B signaling pathway[29–31] in regulating the expression of these adhesion molecules. In the present study, we investigated an alternative mechanism through which Aβ42, and TNF-α regulate the expression of VCAM-1 in BBB endothelial cells while no changes in ICAM-1 expression were observed. Additionally, we investigated the therapeutic potential of SCFA, butyrate in specifically mitigating VCAM-1 expression.

### Signal transducer and activator of transcription 3 (STAT3) and GATA-binding protein 6 (GATA6) expression in AD patients and BBB endothelial models

We investigated the roles of STAT3 and GATA6, two pivotal proteins implicated in the stimulation of TNF-α mediated increases in VCAM-1 expression[32, 33]. Transcription factors play a critical role in modulating endothelial inflammatory responses. The STAT3 transcription factor regulates pro-inflammatory signaling in BBB endothelial cells upon activation by cytokines and oxidative stress[34], the conditions that are present in AD patient brain. Similarly, GATA6 is known to have a direct promoter site on VCAM-1[35, 36] and has been implicated in driving VCAM-1 expression in aortic endothelial cells[37]. Single-nucleus RNA-sequencing data from post-mortem brain samples demonstrated that female AD patients have higher STAT3 levels than age-matched non-demented female controls. In contrast, no changes in male AD patients were observed compared to male or female age-matched non-demented controls (Figure 1A). The RPPA analysis conducted on hCMEC/D3 monolayers exposed to Aβ42 demonstrated a significant increase in pSTAT3 (Tyr 705) and GATA6 levels compared to control cells (Figure 1B).

**Figure 1:**
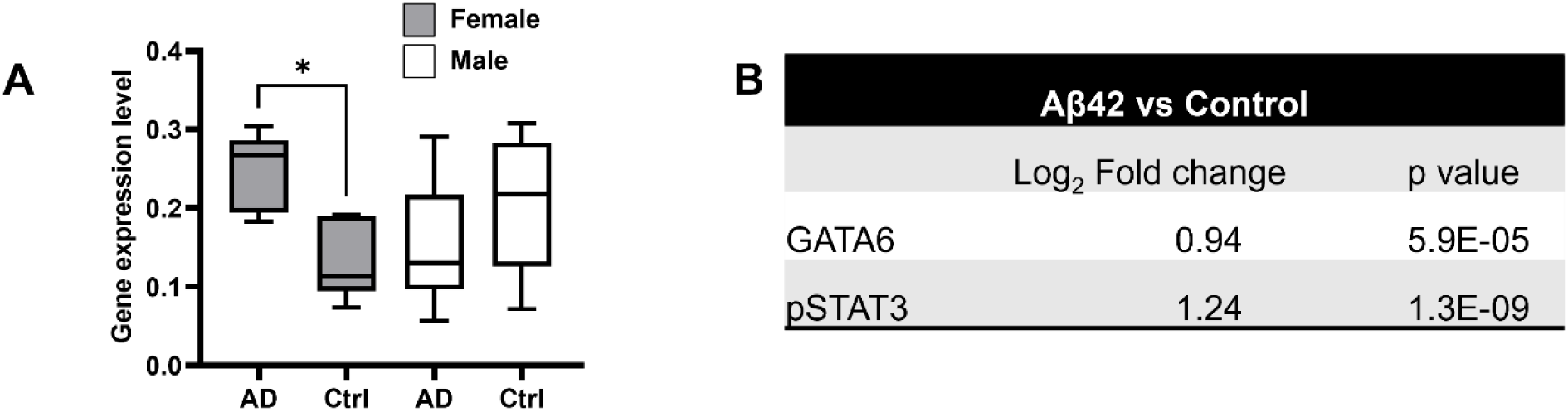
Single nuclei RNA seq and RPPA assay demonstrate changes in the expression of STAT3, pSTAT3 and GATA6. (A) Bar chart depicting gene expression level of STAT3 in Alzheimer’s disease female (AD, gray), normal female (Ctrl, gray), Alzheimer’s disease male (AD, white), and normal male patients (Ctrl, white). Data expressed as mean ± S.D. One-way ANOVA followed by Bonferroni multiple comparison test, *p<0.05. (B) The log2 fold change and p-values demonstrating pSTAT3 and GATA6 protein expression levels in Aβ42-treated cells compared to control hCMEC/D3 cells.

### Butyrate reduces VCAM-1 expression without affecting ICAM-1 expression induced by Aβ42 or TNF-α exposure

Treatment of polarized hCMEC/D3 monolayers with Aβ42 resulted in a two-fold increase in VCAM-1 expression compared to the untreated cells, as demonstrated by confocal microscopy (***p value<0.001, one-way ANOVA). However, pretreating the cell monolayers with butyrate significantly reduced the Aβ42-mediated increase in VCAM-1 expression (*p-value<0.05, one-way ANOVA) (Figure 2A&B). Treatment of hCMEC/D3 monolayers with TNF-α significantly increased the expression of VCAM-1 (∼2-fold, ***p < 0.0001, one-way ANOVA). Previous research has demonstrated that butyrate can attenuate the TNF-α induced upregulation of VCAM-1 expression[25]. In the present study, pretreatment with butyrate effectively reversed TNF-α mediated increase in VCAM-1 expression (*p value< 0.05, one-way ANOVA) (Figure 2C&D). We also investigated the effect of butyrate on ICAM-1 expression in the presence of Aβ42 and TNF-α and demonstrated that Aβ42 increases ICAM-1 expression by ∼ 1.4-fold (***p-value<0.001, One-way ANOVA) and TNF-α increases ICAM-1 expression by ∼1.9-fold (*p < 0.05, one-way ANOVA), as shown by western blot analyses (Figure 2E-H). However, butyrate was unable to significantly reduce Aβ42 (p-value: 0.56, one-way ANOVA) or TNF-α (p= 0.90, one-way ANOVA) mediated increase in ICAM-1 expression (Figure 2E-H).

**Figure 2:**
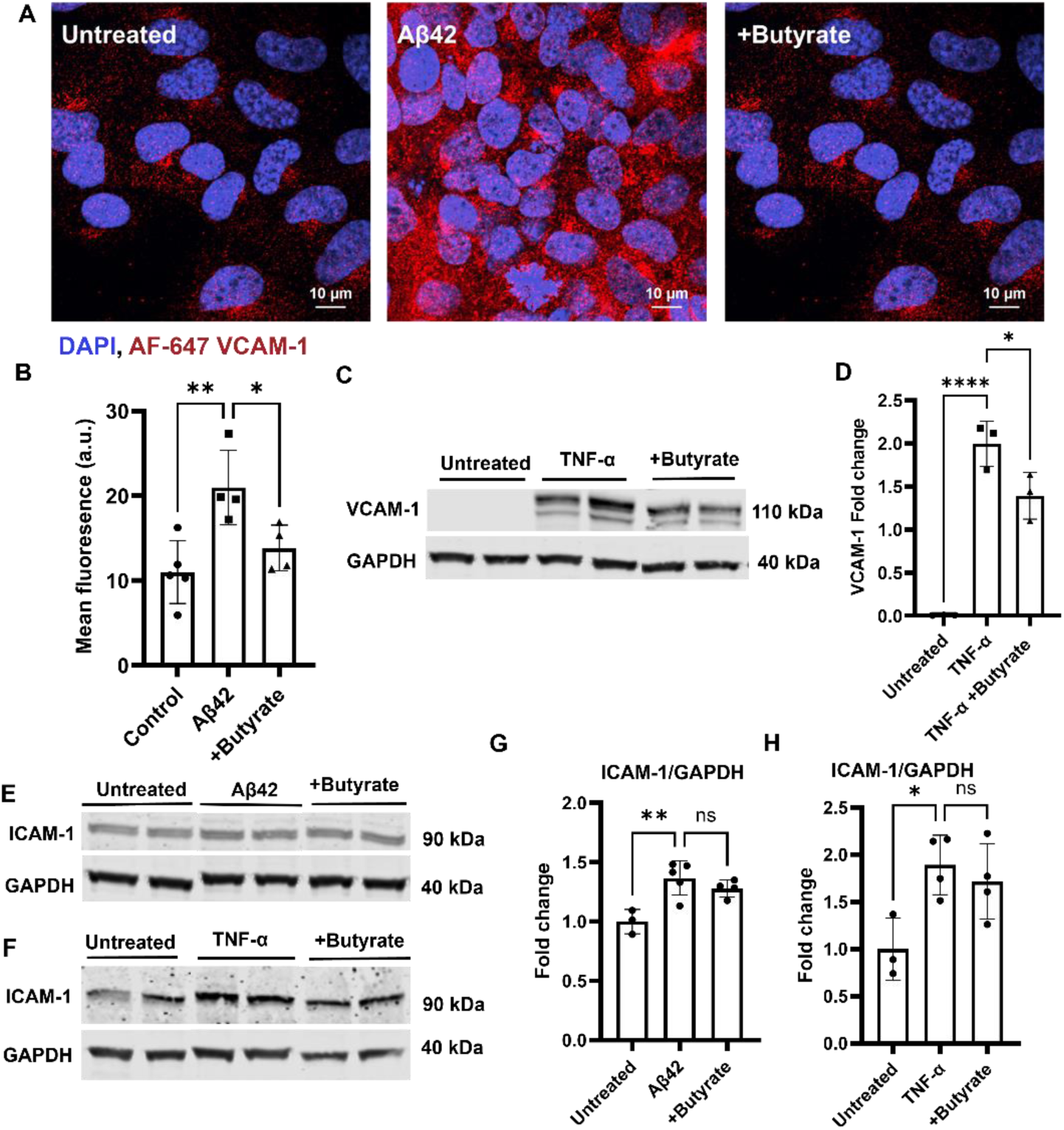
Butyrate mitigates Aβ42 and TNF-α mediated increase in VCAM-1 expression but demonstrated no effect on ICAM-1 expression. Experimental scheme. Polarized hCMEC/D3 monolayers were cultured on 35 mm glass round-bottom dishes or 6-well plates in D3 media. Upon reaching confluence, cells were pretreated with sodium butyrate (10 µM) or left untreated, followed by exposure to Aβ42 (1 μM, 3hrs.) or TNF-α (10 ng/mL, 8 hrs.). (A) Confocal micrographs depict the binding of AF-647 labelled VCAM-1 IgG to the luminal surface of hCMEC/D3 monolayers. The red stain represents AF-647-VCAM-1 IgG, and the nuclei are stained blue (DAPI). Scale bar = 20 μm (B) Bar chart demonstrates quantification of mean fluorescence intensity of confocal micrographs normalized to the cell number. (C) Immunoblots depict VCAM-1 and GAPDH (loading control). (D) The bar chart shows quantification of VCAM-1 immunoblots by densitometry, normalized to GAPDH protein levels. Data expressed as mean ± S.D. One-way ANOVA followed by Bonferroni multiple comparison test. Statistical significance is denoted as follows: *p<0.05, **p<0.01, ****p<0.0001. (E&F) Immunoblots depicting ICAM-1 and GAPDH (loading control) expression in control versus treated hCMEC/D3 monolayers. (G&H) Bar chart shows quantification of immunoblots by densitometry for ICAM-1 normalized to GAPDH protein levels. Data expressed as mean ± S.D. Statistical analysis was performed using one-way ANOVA followed by the Bonferroni multiple comparison test. Statistical significance is denoted as: *p<0.05, **p<0.01

### Butyrate modulates STAT3/GATA6 pathway to decrease VCAM-1 expression

Aβ42 and TNF-α increased VCAM-1 expression in BBB endothelial cells through the activation of GATA6, a well-recognized nuclear transcription factor for VCAM-1[37], as well as STAT3, a major signal transducer. The STAT3 is activated via tyrosine phosphorylation specifically at Y705. Phosphorylation at this site is essential for STAT3 dimerization and subsequent nuclear translocation, which regulates the expression of downstream signaling proteins involved in inflammatory processes[38]. We investigated the effect of butyrate on the expressions of phospho-STAT3 and GATA6 induced by Aβ42 and TNF-α. The hCMEC/D3 monolayers treated with Aβ42 peptide alone showed a significant ∼2-fold increase in phospho-STAT3 expression (*p < 0.05, one-way ANOVA; Figure 3A&B) and a ∼2.2-fold increase in GATA6 expression (**p < 0.001, one-way ANOVA; Figure 3A&C). However, pretreatment with butyrate followed by Aβ42 exposure resulted in a ∼1.7-fold reduction in phospho-STAT3 expression (*p < 0.05, one-way ANOVA; Figure 3A&B) and a ∼1.5-fold reduction in GATA6 expression (*p < 0.05, one-way ANOVA; Figure 3A&C). Similarly, treatment with TNF-α led to a significant ∼2.8-fold increase in phospho-STAT3 expression (**p < 0.0001, one-way ANOVA; Figure 3D&E) and a ∼2.7-fold increase in GATA6 expression (*p < 0.05, one-way ANOVA; Figure 3D&F). Butyrate pretreatment before TNF-α exposure significantly reduced phospho-STAT3 expression by ∼2.3-fold (**p < 0.001, one-way ANOVA) and GATA6 expression by ∼3-fold (***p < 0.0001, one-way ANOVA). These findings demonstrate that butyrate effectively decreases the expression of phospho-STAT3 and GATA6 induced by both Aβ42 and TNF-α in hCMEC/D3 monolayers.

**Figure 3:**
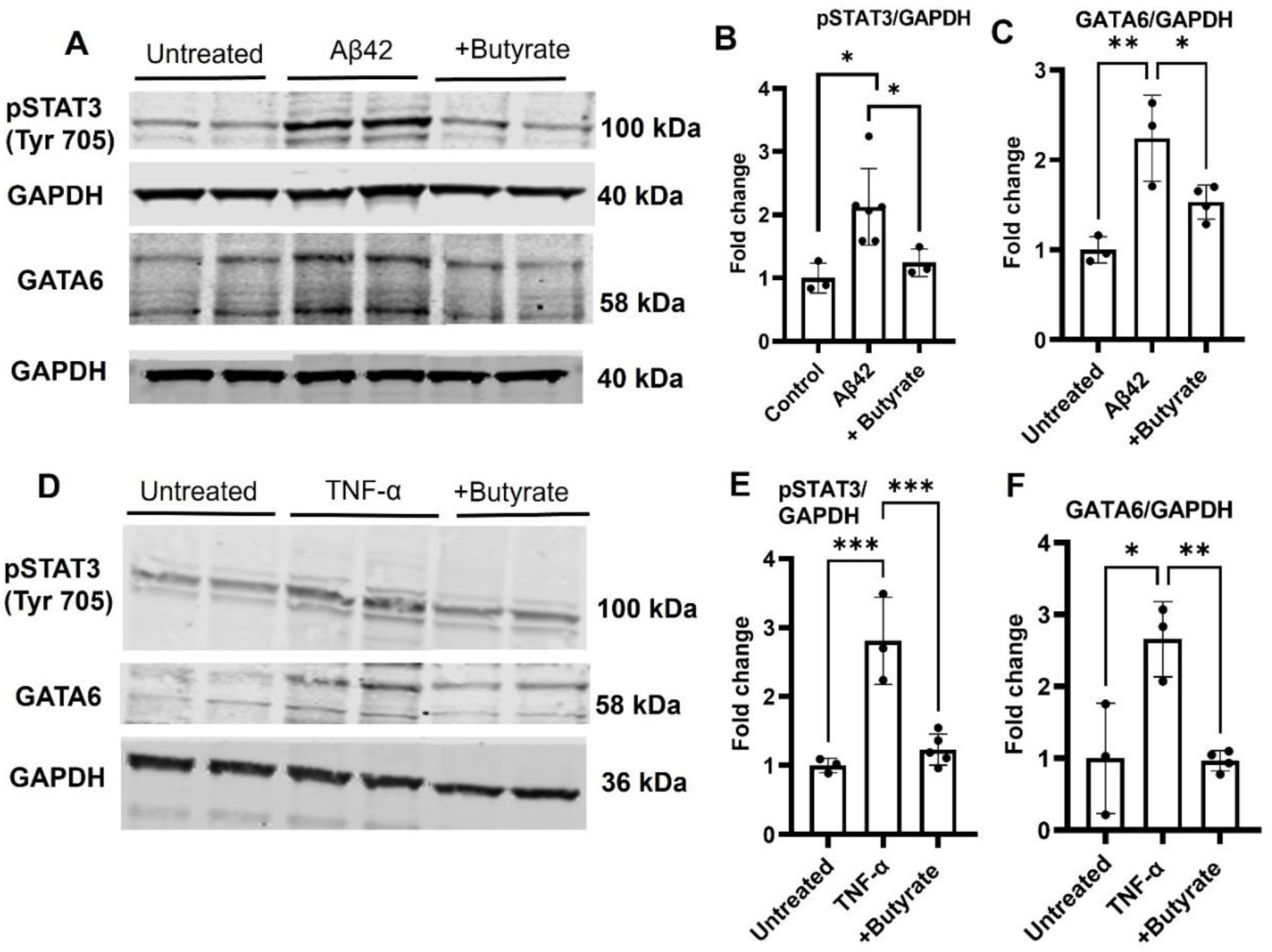
Effect of Aβ42 and TNF-α on STAT3 and GATA6 expression in polarized hCMEC/D3 monolayers with and without butyrate pre-treatment. Experimental scheme. The polarized hCMEC/D3 monolayers were pretreated with 10 μM sodium butyrate or blank buffer, followed by treatment with Aβ42 (1 μM, 3hrs.) or TNF-α (10 ng/mL, 8 hrs.). (A) Immunoblots depicting pSTAT3, GATA6, and GAPDH (loading control) protein levels in hCMEC/D3 lysates upon Aβ42 treatment and (B &C) their densitometry normalized to GAPDH expression. (D) Immunoblots following TNF-α treatment and (E & F) their densitometry normalized to GAPDH expression. Data expressed as mean ± S.D. Statistical analysis was performed using one-way ANOVA followed by Bonferroni multiple comparison test. Statistical significance is denoted as follows: *p<0.05, **p<0.01, ***p<0.001.

### Impact of GATA6 siRNA knockdown on the expression of VCAM-1 and ICAM-1

Pretreatment of hCMEC/D3 monolayers with GATA6 siRNA followed by stimulation with Aβ42 demonstrated a significant reduction (*p value<0.05, unpaired t-test) in VCAM-1 expression, while no changes in ICAM-1 expression were observed (p = 0.18, unpaired t-test) (Figure 4A-D). Similarly, TNF-α exposure to hCMEC/D3 monolayers, in which GATA6 was knocked down with siRNA, resulted in a significant reduction (∼2-fold, **p < 0.001, unpaired t-test) in VCAM-1 expression compared to control cells (Figure 4E&F), whereas no significant effect on ICAM-1 expression was observed. (Figure 4E&H). The knockdown efficiency was confirmed by western blots, showing a significant decrease in GATA6 expression following siRNA treatment (Figure 4I&J)

**Figure 4:**
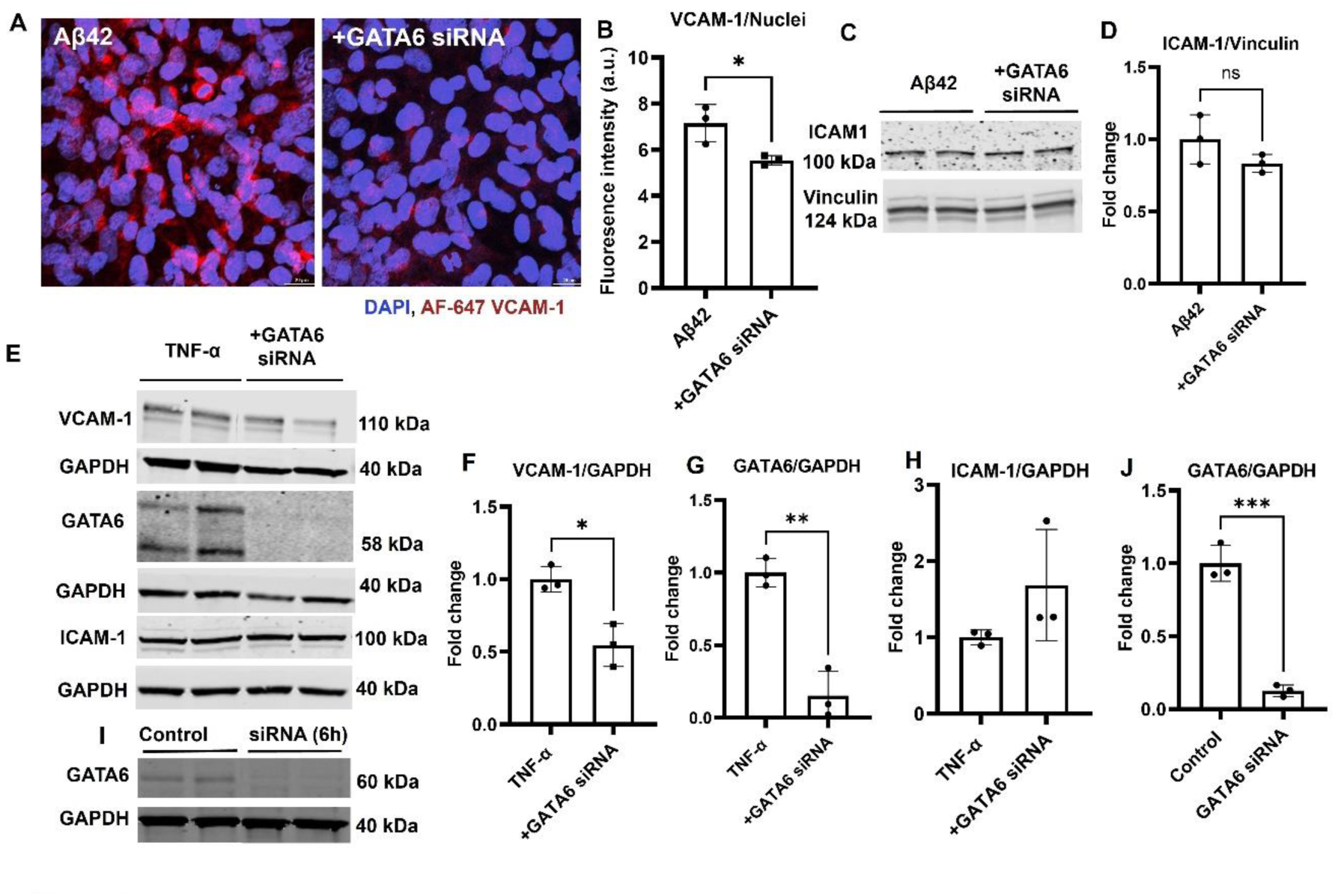
GATA6 knockdown using GATA6 siRNA attenuates Aβ42- and TNF-α induced VCAM-1 expression. Experimental scheme. Polarized hCMEC/D3 monolayers were transfected with GATA6 siRNA for 6 hours or left untreated. Then the treatment media was removed and replaced with fresh D3 media for 24-hour recovery at 37°C. Subsequently, the cells were treated with Aβ42 (1 μM, 3hrs.) or TNF-α (10 ng/mL, 8 hrs.) in low serum (1%) medium to assess VCAM-1 expression. A parallel experiment was conducted with Aβ42 (4 µM, 6 hrs.) or TNF-α (10 ng/mL, 8 hrs.) to evaluate ICAM-1 expression. (A) Confocal micrographs showing Aβ42-induced binding of AF647-VCAM-1 IgG to the luminal surface of hCMEC/D3 monolayers; GATA6 knockdown reverses this effect and reduces VCAM-1 expression. AF-647-VCAM-1 IgG is shown in red and the nuclei are stained with DAPI (blue). Scale bar = 20 μm. (B) Quantification of mean fluorescence intensity from confocal micrographs normalized to cell number. (C) Immunoblot showing ICAM-1 and vinculin (loading control). (D) Bar chart shows quantification of immunoblots by densitometry for ICAM-1 normalized to vinculin protein levels. GATA6 knockdown did not significantly reduce Aβ42-induced ICAM-1 expression (p = 0.18). (E) Immunoblots depict VCAM-1, ICAM-1, GATA6, and GAPDH (loading control). (F-H) Bar chart shows quantification of immunoblots by densitometry for VCAM-1, GATA6, and ICAM-1 normalized to GAPDH protein levels. (I&J) Immunoblot and bar chart demonstrating quantification of magnitude of GATA6 knockdown following GATA6 siRNA treatment normalized to GAPDH loading control. Data are expressed as mean ± S.D. Statistical analysis was performed using one-way ANOVA or unpaired student t-test, as appropriate. Statistical significance is denoted as follows: ns (not significant), *p < 0.05, **p < 0.001, ***p < 0.001.

### The phospho-STAT3 inhibitor cryptothanshinone reduces the expression of VCAM-1

We investigated the effect of cryptothanshinone (CRYPT), a chemical inhibitor of STAT3 activation (phosphorylation of STAT3 at tyrosine 705 site) on VCAM-1 expression induced by Aβ42 and TNF-α. In the hCMEC/D3 monolayers pre-treated with CRYPT, Aβ42 exposure resulted in a significantly lower VCAM-1 expression compared to monolayers treated with Aβ42 alone (****p < 0.0001, one-way ANOVA), as observed via confocal microscopy (Figure 5A&B). Similarly, in hCMEC/D3 monolayers pre-treated with CRYPT followed by TNF-α stimulation, approximately 10-fold reduction in VCAM-1 expression was observed compared to those treated with TNF-α alone (**p < 0.001, one-way ANOVA) (Figure 5C&E). The efficacy of CRYPT in reducing phospho-STAT3 expression was confirmed with 4-hr treatment of hCMEC/D3 monolayers with CRYPT (40 µM), which resulted in a ∼ 2-fold reduction in phospho-STAT3 levels compared to control (*p value<0.05, unpaired t-test) (Figure 5D&F).

**Figure 5:**
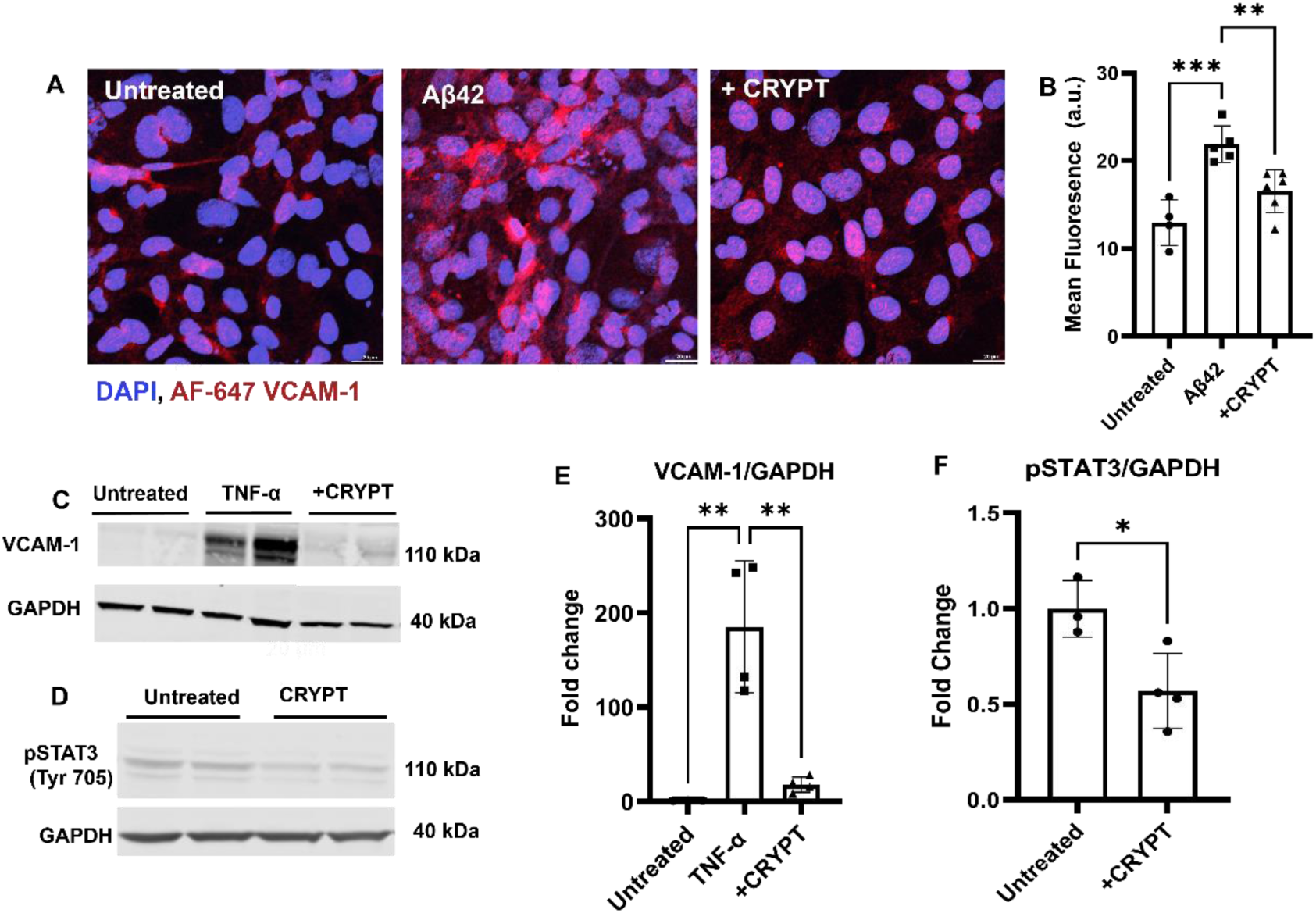
Phospho-STAT3 inhibitor cryptothanshinone (CRYPT) decreases Aβ42 and TNF-α induced VCAM-1 expression. Experimental scheme. Polarized hCMEC/D3 monolayers were pretreated with CRYPT (40 μM, 1 hr) or blank medium, followed by exposure to Aβ42 (1 μM, 3hrs) or TNF-α (10 ng/mL, 8hrs). (A) Confocal micrographs showing AF-647 labeled VCAM-1 IgG binding to the luminal surface of hCMEC/D3 monolayers. AF-647-VCAM-1 IgG is shown in red, and the nuclei are stained with DAPI (blue). Scale bar = 20 μm. (B) Quantification of mean fluorescence intensity from confocal micrographs normalized to cell number (a.u.-arbitrary units). (C&D) Immunoblots showing VCAM-1, phospho-STAT3 and GAPDH (loading control). (E&F) Densitometric quantification of immunoblots for VCAM-1 and phospho-STAT3 normalized to GAPDH. Data presented as mean ± S.D. Statistical analysis was performed using one-way ANOVA or unpaired student t-test, as appropriate. Statistical significance indicated as *p<0.05, **p<0.001.

### STAT3 regulates GATA6 expression

To investigate if GATA6 functions downstream or upstream of STAT3, the effect of phospho-STAT3 inhibition on GATA6 expression was investigated. The hCMEC/D3 cells pre-treated with CRYPT, demonstrated a significant decrease in GATA6 expression compared to those treated with Aβ42 or TNF-α alone (Figure 6A-D) (****p < 0.0001, **p < 0.001, one-way ANOVA).

**Figure 6:**
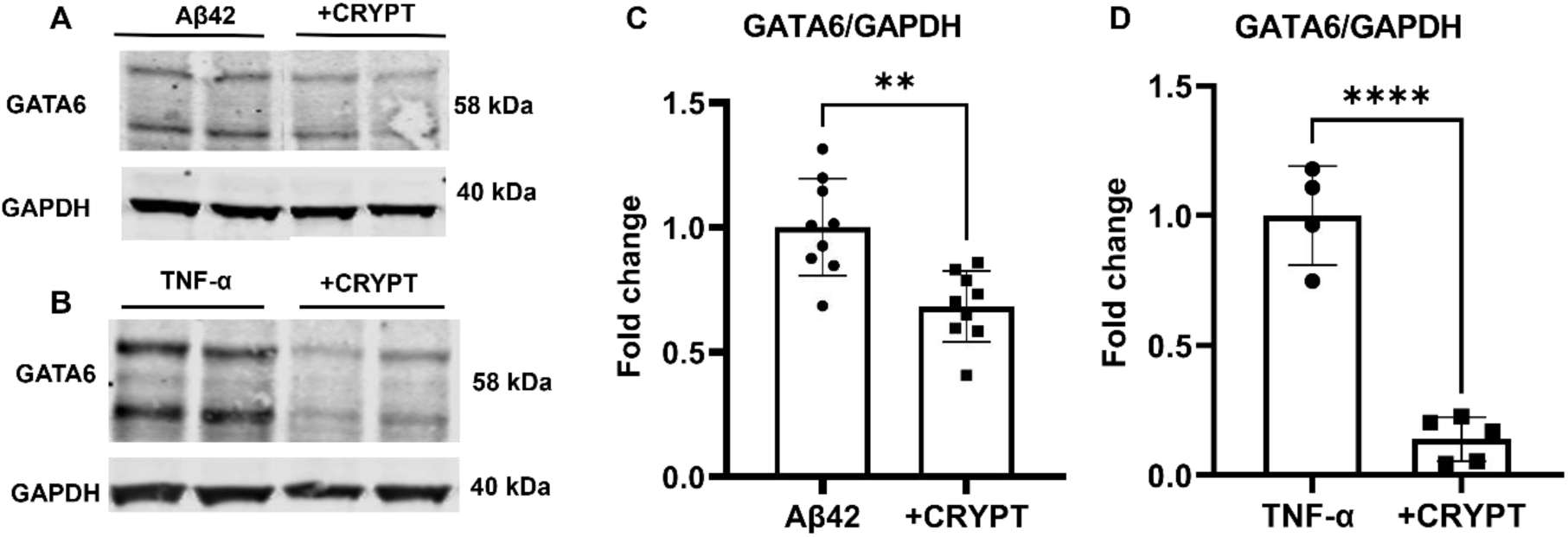
Cryptothanshinone (CRYPT) decreases Aβ42- and TNF-α-mediated increase in GATA6 expression. Experimental scheme. Polarized hCMEC/D3 monolayers were pretreated with CRYPT (40 μM, 1 hr) or blank medium, followed by treatment with Aβ42 (1 μM, 3hrs) or TNF-α (10 ng/mL, 8hrs). (A&B) Immunoblots depict GATA6 and GAPDH (loading control) protein expression. (E&F) Bar charts show quantification of immunoblots by densitometry for GATA6 normalized to GAPDH protein levels, following treatment with Aβ42 (E) or TNF-α (F). Data expressed as mean ± S.D. Statistical analysis was performed using unpaired student t-test. Statistical significance is denoted as follows: ****p<0.00001, **p<0.001.

### Reduced VCAM-1 expression in mice colonized with butyrate-producing bacteria

To assess the impact of butyrate on cerebrovascular inflammation in vivo, we investigated VCAM-1 expression in germ-free mice with targeted knockouts of genes associated with butyrate production. These mice were then subjected to oral gavage with *Clostridium senegalense*, a known butyrate-producing bacterium.

Immunohistochemistry for VCAM-1 demonstrated an increase in VCAM-1 mean fluorescence intensity in the cortex of control germ-free mice compared to those colonized with butyrate-producing bacteria (*p<0.05, paired t-test) (Figure 7A-C).

**Figure 7:**
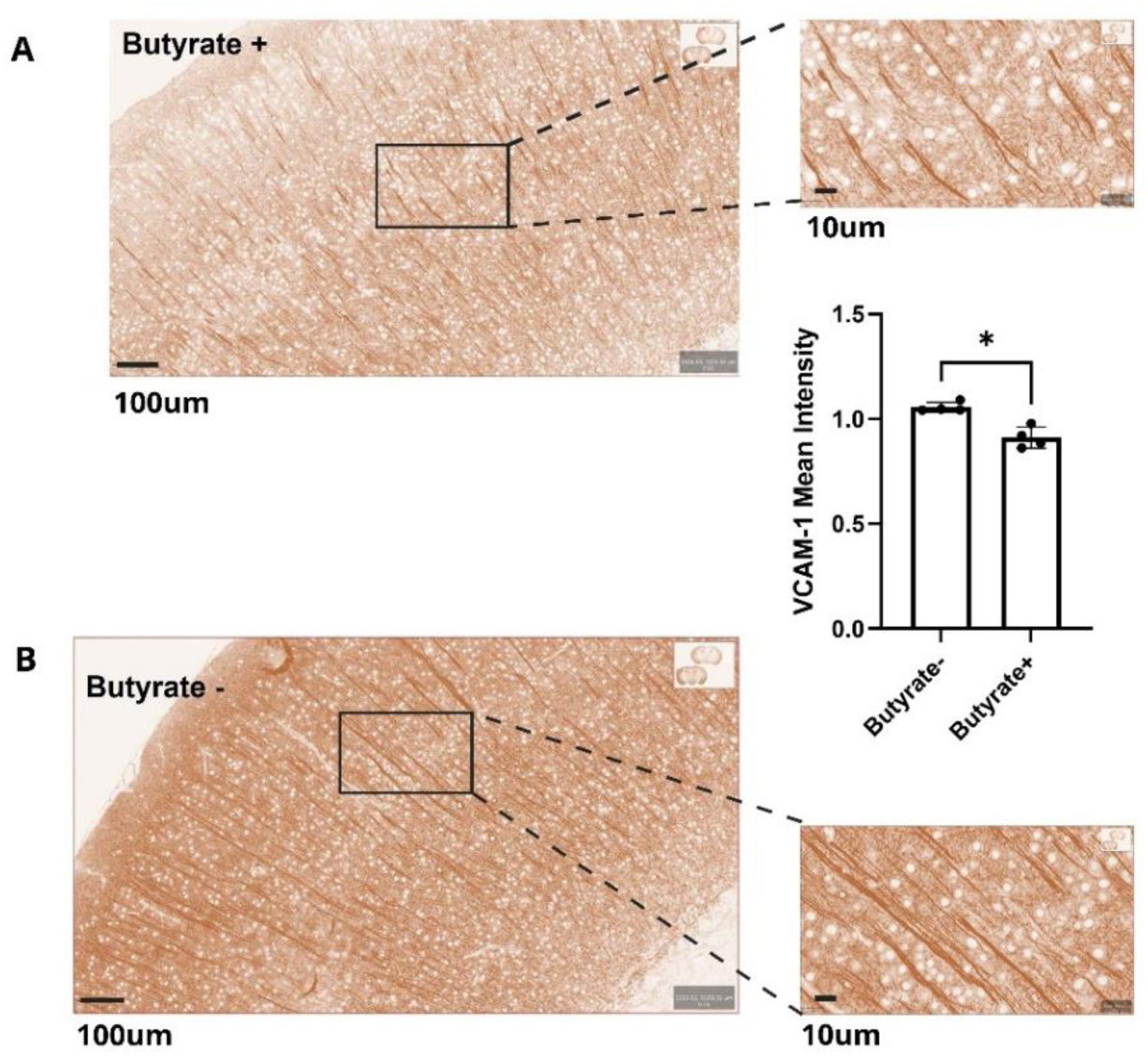
Butyrate reduces VCAM-1 expression in the mouse cortex. (A&B). Representative immunostaining of VCAM-1 performed in cortical brain section from female butyrate (+) and butyrate (-). DAB staining (brown) represents VCAM-1 protein staining intensity, where a stronger DAB staining corresponds to a greater VCAM-1 expression.(C) Bar chart representing quantification of mean DAB intensity in cortical vessels was performed using QuPath from n=4 brain per group. Data represented as mean ±S.D and analyzed using a paired t-test (*p<0.05).

**Figure 8:**
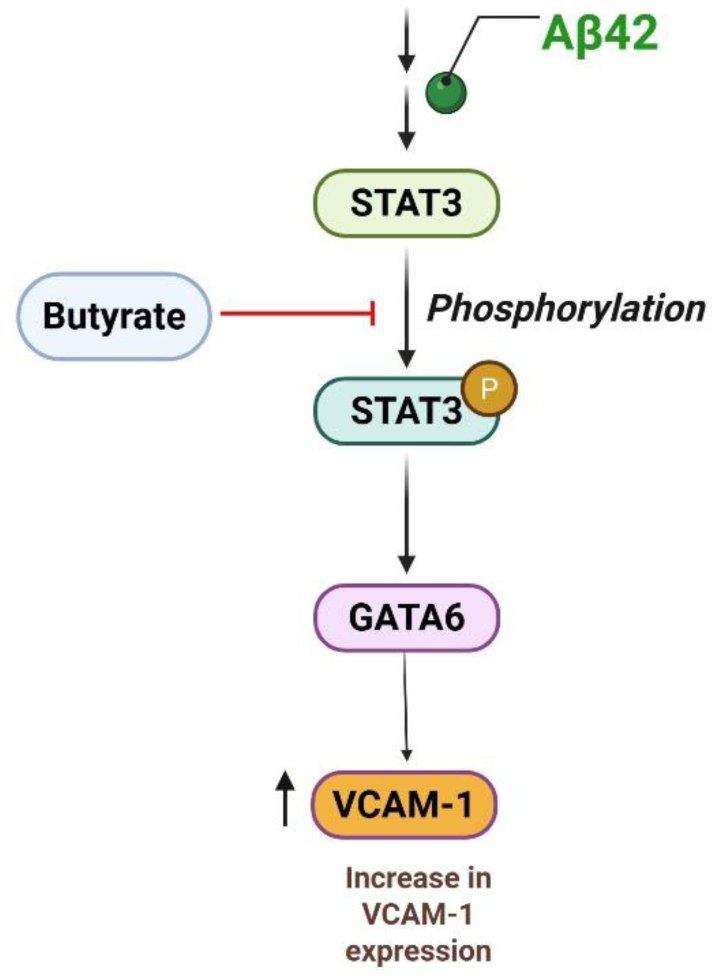
**Butyrate regulates VCAM-1 expression via the STAT3/GATA6 signaling pathway**. Aβ42 stimulates STAT3 activation through phosphorylation at the tyrosine 705 site, which inturn activates GATA6, a transcription factor that induces VCAM-1 expression. Butyrate inhibits this pathway by suppressing STAT3 activation, which downregulates GATA6 expression, leading to a reduction in VCAM-1 expression.

## DISCUSSION

Gut dysbiosis has been implicated in the pathogenesis of AD[39]. Alterations in the gut microbial composition and their metabolic products, particularly SCFAs, are associated with Aβ accumulation in cognitively normal elderly individuals[40–42]. The SCFAs, produced from dietary fiber by gut bacteria, are known to modulate host energy metabolism, inflammation, and immune responses via the gut-brain axis[18, 23, 40].

Among the well-studied SCFAs, butyrate has demonstrated the ability to regulate Aβ burden and improve cognition in preclinical AD mouse models[19, 43]. In addition, the protective effect of butyrate against vascular inflammation has been well documented[44], which suggests that butyrate elicits anti-inflammatory effects that could potentially attenuate cerebrovascular inflammation responses.

Neuroinflammatory changes have been recognized as a crucial contributor to AD progression [45–47], while the role of cerebrovascular inflammation is only recently being recognized[48, 49]. Butyrate was shown to reduce neuroinflammation [50–52]; however, its effect on cerebrovascular inflammation remains only partially understood. This emerging area highlights gaps in our knowledge of how cerebrovascular inflammation contributes to AD progression.

Recent findings by Wyss-Coray and colleagues demonstrated that circulating factors in aged blood induce cerebrovascular inflammation, marked by the upregulation of VCAM-1[53]. Other published reports have shown that elevated levels of TNF-α, a pro-inflammatory cytokine, have been observed in Alzheimer’s patients compared to healthy controls and are also linked to upregulation of adhesion molecules, VCAM-1 and ICAM-1[54, 55]. VCAM-1 is known to sustain chronic inflammation to a greater extent than ICAM-1, as it promotes leukocyte recruitment and immune dysregulation in diseases such as colitis, processes also implicated AD pathogenesis. In contrast, ICAM-1 is primarily involved in acute inflammatory responses and its transient expression may not reflect the sustained cerebrovascular inflammation commonly observed in AD patients [56–58].

Given, the known anti-inflammatory effects of butyrate, we investigated whether butyrate could attenuate cerebrovascular inflammation by modulating VCAM-1 and ICAM-1 expression in brain endothelial cells, following stimulation with soluble Aβ42 or TNF-α. In addition to characterizing the expressions of VCAM-1 and ICAM-1, we also explored the underlying molecular mechanism by which butyrate modulates the expression of these adhesion molecules and identified the involvement of the STAT3-GATA6 signaling pathway.

Previous studies have reported that butyrate pretreatment reduces VCAM-1 expression following TNF-α stimulation in human umbilical vascular endothelial cells (HUVECs)[25, 44], while no changes were observed in ICAM-1 expression[25]. Specifically, pre-treatment of HUVECs with butyrate at a supraphysiological concentration of 10 mM decreased VCAM-1 expression stimulated by TNF-α. Similarly, Zapolska-Downer et al. reported that butyrate attenuated TNF-α and IL-β induced VCAM-1 expression to a greater extent than ICAM-1 expression in HUVEC cell monolayers[59]. However, the mechanisms underlying this differential regulation of VCAM-1 and ICAM-1 expressions were not investigated. Furthermore, the effect of Aβ42 on VCAM-1 expression in the presence of butyrate has not been investigated in the BBB endothelium.

In the current study, we showed that polarized hCMEC/D3 monolayers treated with Aβ42 peptide or TNF-α increases both VCAM-1 and ICAM-1 expression. However, pretreatment with butyrate reduced the Aβ42 and TNF-α mediated increase in VCAM-1 expression without impacting ICAM-1 expression, suggesting butyrate’s selective anti-inflammatory effect. To investigate the molecular mechanisms underlying this selective regulation, we examined transcriptional mediators that modulate adhesion molecule expression. Transcription factors such as nuclear factor kappa-B (NF-*κ*B), interferon regulatory factor-1 (IRF-1), GATA6, and activator protein-1 (AP-1) are recognized to regulate the expression of VCAM-1 and ICAM-1. Treatment of various cell lines with chemical inhibitors and genetic silencing of these transcription factors using siRNA were shown to reduce VCAM-1 and ICAM-1 expression[33, 60, 61]. The VCAM-1 promoter contains binding sites for NF-κB, AP-1, IRF-1, and GATA6[62, 63], while the ICAM-1 promoter has functional binding sites for NF-κB and AP-1[37], but lacks IRF-1 and GATA6 binding sites. This difference in promoter sites may explain the differential regulation of VCAM-1 and ICAM-1 to butyrate. Previous studies have demonstrated that GATA6 regulate VCAM-1 and not ICAM-1 expression, which suggests butyrate potentially may act through this pathway to selectively reduce VCAM-1 expression[64, 65].

Butyrate is widely recognized as an inhibitor of class I and II histone deacetylases (HDAC 1/2). Chang et al. demonstrated that butyrate exhibits anti-inflammatory effects by inhibiting HDACs[66]. Based on these findings, we hypothesized that butyrate’s differential effects on VCAM-1 and ICAM-1 expression may be mediated by its HDAC1/2 inhibitory activity. Hu et al. 2021 have demonstrated that treatment with romidepsin, a selective HDAC1/2 inhibitor, decreased TNF-α mediated increase in VCAM-1 expression without affecting ICAM-1 expression in aortic endothelial cells [37]. Supporting this hypothesis, Zhou et al. reported that knocking down GATA6 with siRNA reduced TNF-α-mediated increase in VCAM-1 expression in human aortic endothelial cell line[32]. Similarly, in our studies, knockdown of GATA6 in hCMEC/D3 monolayers using siRNA, followed by stimulation with Aβ42 or TNF-α, resulted in a reduced VCAM-1 expression, while ICAM-1 expression remained unchanged. Additionally, our studies have shown that butyrate pre-treatment reduced Aβ42 and TNF-α induced GATA6 expression in hCMEC/D3 monolayers. Taken together, these results highlight the selective role of GATA6 in regulating VCAM-1 expression, while not affecting ICAM-1 expression, and suggest that butyrate exerts its anti-inflammatory effects through the GATA6 axis.

Our study further identified STAT3 as a non-histone substrate of HDAC 1/2 that regulates VCAM-1 expression via GATA6. STAT3 is a cytokine-inducible transcription factor activated by phosphorylation at the tyrosine 705 site [67] [68]. Previous studies have implicated STAT3 in TNF-α mediated upregulation of VCAM-1 expression [32], but this pathway has not been explored in the BBB endothelium. We have shown that exposure of hCMEC/D3 monolayers to Aβ42 or TNF-α promotes STAT3 phosphorylation at tyrosine 705, and this effect was attenuated by butyrate pretreatment. Pharmacological inhibition of STAT3 with cryptothanshinone, significantly reduced Aβ42 and TNF-α induced VCAM-1 expression, confirming STAT3 as an upstream regulator in this pathway.

Finally, we examined the interaction between STAT3 and GATA6 in regulating VCAM-1 expression. Zhou et al. identified multiple STAT3 binding sites within the GATA6 promoter[32], suggesting that STAT3 directly regulates GATA6 expression. Consistent with these findings, our data demonstrated that inhibition of STAT3 with cryptothanshinone reduced Aβ42 and TNF-α induced GATA6 expression. Together, these findings support the presence of a signaling cascade in which Aβ42 or TNF-α activates STAT3 phosphorylation, which in turn upregulates GATA6, ultimately driving VCAM-1 expression in BBB endothelial cells. Butyrate disrupts this pathway by suppressing STAT3 activation and thereby attenuating GATA6-mediated VCAM-1 upregulation.

To extend our findings beyond the in vitro setting, we investigated the in vivo relevance of butyrate in modulating cerebrovascular inflammation. To our knowledge, this study represents the first application of a microbial colonization model using a butyrate-deficient mutant engineered by targeted deletion of KamD and crotonase genes, to assess the role of microbiota-derived butyrate in regulating VCAM-1 expression. This experimental approach facilitated the assessment of the effects of endogenous butyrate depletion on VCAM-1 expression while minimizing confounding factors such as antibiotic-induced dysbiosis and broader gut microbial disruption.

Histological analysis revealed that mice colonized with the butyrate-deficient strain (butyrate-) exhibited higher mean intensity of DAB staining, which translates to a greater increase in VCAM-1 expression compared to the butyrate-sufficient group (butyrate+). Additionally, the butyrate-group showed a greater density of VCAM-1 expressing blood vessels, marked by the intense and widespread brown staining along vessel-like structures, particularly within cortical regions, indicating increased endothelial activation. The pronounced VCAM-1 signal seen in these butyrate-mice underscores the pro-inflammatory environment associated with butyrate depletion. In contrast, the butyrate+ condition displayed a markedly reduced VCAM-1 signal, with fewer and more faintly stained vascular elements, indicating a reduction in activation of pro-inflammatory signals in endothelial cells. These findings demonstrate that endogenous butyrate production is essential to maintain cerebrovascular homeostasis by reducing BBB endothelial inflammation.

In summary, our study establishes that butyrate mitigates Aβ42 and TNF-α-induced cerebrovascular inflammation by suppressing VCAM-1 expression in BBB endothelial cells through the inhibition of the STAT3-GATA6 signaling axis. These findings highlight butyrate’s therapeutic potential to target cerebrovascular inflammation, widely believed as a key contributor to AD pathogenesis.

## MATERIAL AND METHODS

### Animals

Female BL6 mice at around 5 weeks of age were used. The C. senegalense DSM 25507 control and C. senegalense DSM 25507 mutant were initially cultured on pre-reduced TSA plate with 15 ug/mL thiamphenicol and 10 ug/mL erythromycin, from which single colonies were inoculated into liquid BHI medium. All liquid culture for gavaging did not include any antibiotics to avoid effects on subsequent in vivo studies. Mice were colonized with either *Clostridium senegalense* DSM 25507 WT (butyrate-producing), or the double-KO mutant (no butyrate-producing), which was knocked out of *KamD* (gene in the lysine pathway for butyrate production) and *crotonase* (gene in the acetyl-CoA pathway for butyrate production) genes. The colonization was done by an oral gavage of 300 uL of overnight anaerobic culture of the bacteria in pre-reduced brain heart infusion medium acquired. The mice were administered the oral gavage at 3.5-4.5 weeks of age and maintained for 12 days before sampling. Animals were then maintained in the Tecniplast IsoCage system for 2 weeks. The IACUC#A00005635 approved by Mayo for animal housing and maintenance setup was followed.

### Cell culture

The immortalized human cerebral microvascular endothelial cell (hCMEC/D3) line was generously provided by P-O Couraud of Institut Cochin, France. The endothelial cells were cultured following previously established protocols[69] in endothelial basal media (Sigma-Aldrich,St. Louis, MO, supplemented with various reagents that support cell growth. These reagents include a chemically defined lipid concentrate (1% v/v, ThermoFisher Scientific, Waltham, MA), penicillin-streptomycin (1% v/v, Sigma-Aldrich, St. Louis, MO), recombinant human fibroblast growth factor (1 ng/mL, PeproTech, Rocky Hill, NJ), hydrocortisone (1.4 μM, Sigma-Aldrich, St. Louis, MO), ascorbic acid (5 μg/mL, Sigma-Aldrich), HEPES (10 mM), and fetal bovine serum (1% or 5%, Atlanta Biologicals, Flowery Branch, GA). This endothelial basal media, along with the supplements, is referred to as D3 media. The polarized hCMEC/D3 monolayers were cultured in collagen-coated (Corning, MA) 35 mm glass coverslip bottomed dishes (Mattek, MA) or 6-well plates (Corning Costar, MA) at 5% CO2 and 37°C in D3 media with 5% serum. The medium was changed every 2 days until the cells reached confluence (∼4-5 days). The cells from passages 34–35 was used for the experiments.

### Western blotting

The hCMEC/D3 monolayers were pretreated with sodium butyrate (10 µM) under specific conditions, 21 hours for the evaluation of VCAM-1 expression and 18 hours for the assessment of ICAM-1 expression. Following this pretreatment, cells were exposed to Aβ42 (1 µM, 3 hours) or TNF-α (10 ng/mL, 8 hours) for VCAM-1 analysis. In the case of ICAM-1, the cells were stimulated with Aβ42 (4 µM, 6 hours) or TNF-α (10 ng/mL, 8 hours). The cells were washed three times with ice-cold PBS and lysed using radioimmunoprecipitation assay buffer containing protease and phosphatase inhibitors (Sigma-Aldrich, St. Louis, MO). The total protein concentration of the whole-cell lysates was determined using the bicinchoninic acid assay (Pierce, Waltham, MA). Whole-cell lysates containing 20-30 µg of protein were prepared and loaded on 6%-12% polyacrylamide gels and the proteins were resolved by SDS-PAGE under reducing conditions (Bio-Rad Laboratories, Hercules, CA). The gels were electrophoretically transferred to a nitrocellulose membrane (0.2 µm pore size). Subsequently, the nitrocellulose membrane was blocked for 1 hour at room temperature using 5% w/v milk protein in tris-buffered saline with 0.1% Tween20 detergent (TBST). Primary antibodies targeting specific proteins [phospho-STAT3 (1:1000), GATA6 (1:500), VCAM-1 (1:1000), ICAM-1 (1:250), and protein loading controls [GAPDH (1:1000) or vinculin (1:1000)] were incubated overnight at 4 °C. After the overnight incubation with the primary antibody, the nitrocellulose membrane was washed with TBST to remove any unbound antibody. Then the membrane was incubated with a secondary antibody conjugated to a near-infrared (800 nm) IR-dye at a 1:2000 dilution for one hour at room temperature.

The protein bands were visualized using an Odyssey Licor instrument (Odyssey CLx; LI-COR Inc.) and quantified using Image Studio Lite 5.2 software. The intensity of the bands on the nitrocellulose membrane were normalized to the intensity of the GAPDH or vinculin band, which served as the loading control.

### Aβ peptide film preparation and Reconstitution

The unlabeled Aβ peptides were procured from Aapptec (Louisville, KY) and were assayed to be of 95% purity as determined by high performance liquid chromatography-mass spectrometry. The monomeric solutions of Aβ peptides were prepared according to the procedure previously published by Klein[70]

### Reverse phase protein array assay (RPPA assay)

The hCMEC/D3 cells were seeded on 24 mm Transwells^®^ (100,000 cells per well) and grown as polarized monolayers in D3 media. After 7 days, when the polarized monolayers are expected to be fully formed, the Transwells^®^ were switched to low serum D3 media (1% FBS) a day before the experiment. The cells were treated with or without 1 μM of Aβ42 for 1 hour on the luminal side of the Transwell^®^ at 37 °C. Following the incubation, the cells were washed twice with Hank’s balanced salt solution buffer containing protease and phosphatase inhibitors. Following the wash steps, radio immunoprecipitation assay buffer was added to each Transwell^®^ to lyse the cells and the cells were detached using a cell scrapper. The samples were centrifuged at 10,000 rpm for 10 minutes at 4 °C, the pellet was discarded, and the supernatant was stored at -20 °C. The frozen samples were shipped for analysis at the RPPA core facility in MD Anderson Cancer Center, Houston, TX. The RPPA analysis was conducted according to protocols optimized by the core facility. Briefly, the lysates were diluted and blotted on nitrocellulose slides. The slides were then probed with 305 antibodies using a tyramide-based signal amplification approach and visualized by a DAB colorimetric reaction.

Slides were scanned and density was quantified by an Array-Pro analyzer. Relative protein levels were determined by interpolation using super curve, which is a software that MD Anderson uses to generate the RPPA data[71, 72]. The relative protein levels were presented as raw log2 transformed data, which was centered using the sample means. A contrast model was constructed using limma (v 3.46.0) between the control vehicle and Aβ42. Pairwise post hoc analyses were conducted for antibodies that exhibited significance (α = 0.05), confirming the initially observed differences in the contrast model. One way ANOVA was then performed to test the significance of differences between the control group and the Aβ42 treated group.

### Single nucleus RNA sequencing analysis in Alzheimer’s patients

Single-nucleus RNA-sequencing data from post-mortem brain samples of 24 patients in the ROSMAP cohort were downloaded from Synapse (syn16780177). This cohort comprised 12 female patients and 12 male patients, with each group including 6 individuals diagnosed with AD and 6 who were non-cognitively impaired. A total of 162,767 single nuclei were processed and clustered into 25 cell type-specific clusters, as annotated by the data provider. The endothelial cluster (N=1,969) was selected for further analysis. Differential expression (DE) analysis was performed using the non-parametric Wilcoxon rank-sum test (Seurat Find Markers) to compare AD and non-cognitively impaired patients, separately for males and females. Genes with an adjusted p-value < 0.05 were considered significantly differentially expressed

### GATA6 knockdown

GATA6 knockdown was performed in hCMEC/D3 monolayers that are 60%-70% confluent. The knockdown was performed with GATA6 siRNA (Santa Cruz, TX), RNAi max (Invitrogen, CA), and Opti-MEM (Gibco, NY) in reduced serum medium. Following transfection, the cells were allowed to recover for 24 hours and then lysed to conduct western blot analysis.

### Immunocytochemistry and cellular imaging

Polarized hCMEC/D3 cell monolayers were cultured on 35 mm round bottom glass dishes in D3 media containing 5% FBS. The cells were transitioned to D3 medium containing 1% FBS, a day before the experiment. The cells were pre-treated with either butyrate (10 µM), cryptothanshinone (40 µM), or blank D3 media for 21 hours.

Subsequently, Aβ42 (1 µM) was added to the media and co-incubated with or without butyrate or cryptothanshinone for 3 hours at 37°C, while the control samples were maintained in blank D3 medium.

After the treatment period, the cells were washed three times with ice cold PBS, fixed with 4% paraformaldehyde, and blocked using 10% goat serum (Invitrogen, CA). Then the hCMEC/D3 monolayers were incubated with unlabeled VCAM-1 antibody (Abcam, Cambridge, UK) at a 1:250 dilution and incubated overnight at 4°C followed by incubation with AF-647 secondary antibody (1:1000) for 1 hr. at room temperature. Finally, the hCMEC/D3 monolayers were washed with ice-cold PBS and treated with 3 µM DAPI (Invitrogen) solution for 10 minutes at room temperature. The monolayers were subsequently washed with PBS, mounted with Prolong Gold antifade mounting medium (Invitrogen), and imaged with Nikon Confocal A1Rsi NSIM confocal microscope using a 60X,1.4NA objective (University of Minnesota Imaging Center). Data was acquired using NIS Elements AR software (v.5.6, Nikon Instruments USA, Melville, NY). The images were quantified using Fiji software where the mean fluorescence intensity of each image was quantified and normalized to the cell number, determined using the cell counter function in Fiji.

### Immunohistochemistry

The BL6 mice brains were fixed in formalin and embedded in paraffin and sectioned into 5-μm-thick brain slices. The tissue sections on the slides were melted for 1 h at 60 °C, deparaffinized in xylene and ethanol, and rehydrated in TBS. Antigen retrieval was performed by placing the slides in 10 mM citrate buffer (pH 6.0) and heating them at 95 °C for 45 minutes, followed by cooling for 30 minutes and washing with PBS.

Endogenous peroxidase activity in the sections was blocked in 5% hydrogen peroxide in TBS for 10 minutes. Sections were washed with TBS and blocked for 1 hour with 10% v/v serum in TBS. Then the sections were incubated with the rabbit anti-VCAM-1 polyclonal antibody (1:4000 dilution) for VCAM-1 detection in a humid chamber overnight at 4 °C. The sections were rinsed and incubated for 2 h at room temperature with peroxidase-conjugated secondary antibodies. Subsequently, they were rinsed with PBS and stained using the DAB Plus substrate staining system as per the manufacturer’s instructions. All tissue sections were counterstained with hematoxylin and mounted. The brain sections were scanned in ribbons with a Huron TissueScope LE brightfield imaging system (Huron Digital Pathology) at 40X magnification (0.75 NA) with a pixel resolution of 0.25 um. Images were quantified using QuPath (version 0.5.1). For each mouse, 50-70 regions of interest (ROIs) were manually selected using the wand tool to isolate VCAM-1 positive vessels within the cortex. The mean pixel intensity at a resolution of 0.22 µm was obtained from each ROI and the average of all ROIs was calculated to yield a single mean value per mouse brain. Each slide included both experimental groups (n=4) to control for staining variability. Statistical analysis was conducted using paired t-test in GraphPad Prism (version 10.2.3).

## Acknowledgments

### Funding

National Institute of Health/National Institute of Neurological Disorders and Stroke R01NS125437 (KKK)

National Institute of Health DK114007 (PKC).

Imaging work was supported by the resources at the University of Minnesota University Imaging Centers (UIC)-SCR_020997.

We are grateful for a grant-in-aid (#324930) from the University of Minnesota for the purchase of the LI-COR Odyssey CLx Infrared imaging system that supported western blot imaging.

Immunohistochemistry was performed at the Histology core, University of Minnesota.

### Author contributions

Conceptualization: VSS, KKK

Methodology: VSS, VV, KKK, YX

Investigation: VSS, VV, KKK, YX

Visualization: VSS, VV, XT, KKK

Funding acquisition: KKK, PCK

Supervision: KKK, PCK, KRK

Writing – original draft: VSS, KKK

Writing – review & editing: VSS, VV, YX, XT, PCK, KKK

## Data and materials availability

All data are available in the main text or the supplementary materials.

## Acknowledgements

Figures were created on Biorender.com

## Data Availability Statement

The authors declare that all the data supporting the findings of this study are contained within the paper.

